# Thermospermine is an evolutionarily ancestral phytohormone required for organ development and stress responses in the basal land plant *Marchantia polymorpha*

**DOI:** 10.1101/2023.07.13.548936

**Authors:** Takuya Furumoto, Shohei Yamaoka, Takayuki Kohchi, Hiroyasu Motose, Taku Takahashi

## Abstract

Thermospermine, a structural isomer of spermine, suppresses auxin-inducible xylem differentiation, whereas spermine is implicated in stress responses in angiosperms. Thermospermine synthase ACAULIS5 (ACL5) is well conserved from algae to land plants, but its physiological function remains elusive in non-vascular plants. Here we focused on Mp*ACL5*, a gene in the liverwort *Marchantia polymorpha*, which rescued the dwarf phenotype of the *acl5* mutant of *Arabidopsis*. In the Mp*acl5* mutants generated by genome editing, growth of the vegetative organ, thallus, and the sexual reproductive organ, gametangiophore, was severely retarded. The mutant gametangiophore exhibited remarkable morphological defects such as short stalks, fasciation, and indeterminate growth; it was formed as a fusion of two gametangiophores and a new gametangiophore was often initiated from the old one. Furthermore, Mp*acl5* was shown to be hypersensitive to heat and salt stresses. Given the absence of spermine in liverworts including *M. polymorpha*, these results reveal that thermospermine has a dual primordial function in organ development and stress responses in the basal land plant lineage, the latter of which may have eventually been assigned to spermine during the land plant evolution.

## Introduction

Polyamines are low-molecular-weight aliphatic compounds with multiple amino groups. They are involved in various biological activities including stabilization of nucleic acids, mRNA translation, and modulation of protein functions (Igarashi and Kashiwagi 2010, Pegg 2016). The diamine putrescine and the triamine spermidine are ubiquitously present and essential in all organisms. The tetraamine spermine is generally found in animals and fungi but not always detected in bacteria and plants (Takano et al. 2012). There are to date no genes identified in mosses and ferns for spermine synthase, an aminopropyl transferase that mediates the synthesis of spermine from spermidine and an aminopropyl moiety provided by decarboxylated S-adenosylmethionine. In *Arabidopsis*, a loss-of-function mutant of a single gene for spermine synthase shows no morphological phenotype under normal growth conditions (Imai et al. 2004). However, there are increasing evidence that spermine is implicated in the response to biotic and abiotic stresses in angiosperms (Tiburcio et al. 2014).

Thermospermine is a structural isomer of spermine and was first found in an extra thermophilic bacterium (Oshima et al. 1979). In contrast to spermine, thermospermine is widespread in the plant kingdom. The gene encoding thermospermine synthase, which was identified from its mutant *acaulis5* (*acl5*) in *Arabidopsis* (Hanzawa et al. 1997) and named *ACL5*, may have been acquired early in the plant evolution by horizontal gene transfer from bacteria (Minguet et al. 2008). *ACL5* is exclusively expressed in xylem precursor cells and involved in suppressing excessive xylem formation (Clay and Nelson 2005). The excess xylem phenotype associated with the dwarf growth of *acl5* is partially restored by exogenous thermospermine (Kakehi et al. 2008). Isolation and analysis of *suppressor of acl5* (*sac*) mutants revealed that thermospermine enhances mRNA translation of *SAC51*, which encodes a basic helix-loop-helix (bHLH) protein and plays a role in suppressing xylem differentiation (Imai et al. 2006). Thermospermine may release the *SAC51* mRNA from an inhibitory effect of its own conserved upstream open reading flame (uORF) on the main ORF translation, although the precise mode of action remains unknown (Imai et al. 2006, 2008, Kakehi et al. 2015, Ishitsuka et al. 2019). Furthermore, a series of genetic analysis have shown the involvement of the *SAC51* family in thermospermine-mediated negative feedback regulation of auxin-inducible xylem differentiation (Vera-Sirera et al. 2015, Katayama et al. 2015). Auxin promotes expression of *TARGET OF MONOPTEROS 5* (*TMO5*) encoding a bHLH transcription factor, which form a heterodimer with another bHLH transcription factor LONESOME HIGHWAY (LHW). TMO5-LHW complex promotes xylem proliferation and induces expression of *ACL5* and *SACL3*, a member of the *SAC51* family whose product in turn competes with TMO5 for the binding with LHW to interfere with excess xylem formation (Vera-Sirera et al. 2015, Katayama et al. 2015).

Bryophytes including liverworts and mosses do not develop vascular systems but have genes highly homologous to *ACL5*, which suggests as-yet-unidentified fundamental function of thermospermine in the basal land plants (Solé-Gil et al. 2019). Thus, we focused on the single gene in the liverwort, *Marchantia polymorpha*, Mp*ACL5*. In this study, we have generated a knockout mutant of Mp*ACL5.* Characterization of the mutant revealed that thermospermine is critically required for both organ development and stress responses in this liverwort.

## Results

### Mp*ACL5* is preferentially expressed in meristematic regions

The Mp*ACL5* gene consists of nine exons and eight introns whose positions are well conserved in land plants and encodes a protein of 340 amino acid residues with a molecular mass of 37.7 kDa, which shows 63% identity to *Arabidopsis* ACL5 (Solé-Gil et al. 2019, Supplementary Fig. S1). The MpACL5 protein has been shown to function as thermospermine synthase in yeast cells (Solé-Gil et al. 2019). We confirmed that the recombinant MpACL5 expressed in *Escherichia coli* can also synthesize thermospermine (Supplementary Fig. S2). To determine whether MpACL5 functions as thermospermine synthase *in planta*, we generated transgenic *Arabidopsis* lines carrying the Mp*ACL5* cDNA driven by the constitutive cauliflower mosaic virus (CaMV) 35S promoter in the *acl5* mutant background and confirmed that the dwarf phenotype of *acl5* was rescued (Fig. 1). Reverse transcriptase-PCR experiments revealed that the Mp*ACL5* transcript was detected in all organs tested including thallus, gemma cup, and gametangiophore (Fig. 2A). We also generated transgenic *M. polymorpha* lines expressing Citrine fluorescent protein with a nuclear localization signal (*Citrine-NLS*) under the control of the Mp*ACL5* promoter. Strong Citrine-NLS fluorescence was observed in meristematic apical notches of the thallus margin (Fig. 2B).

**Fig. 1.**
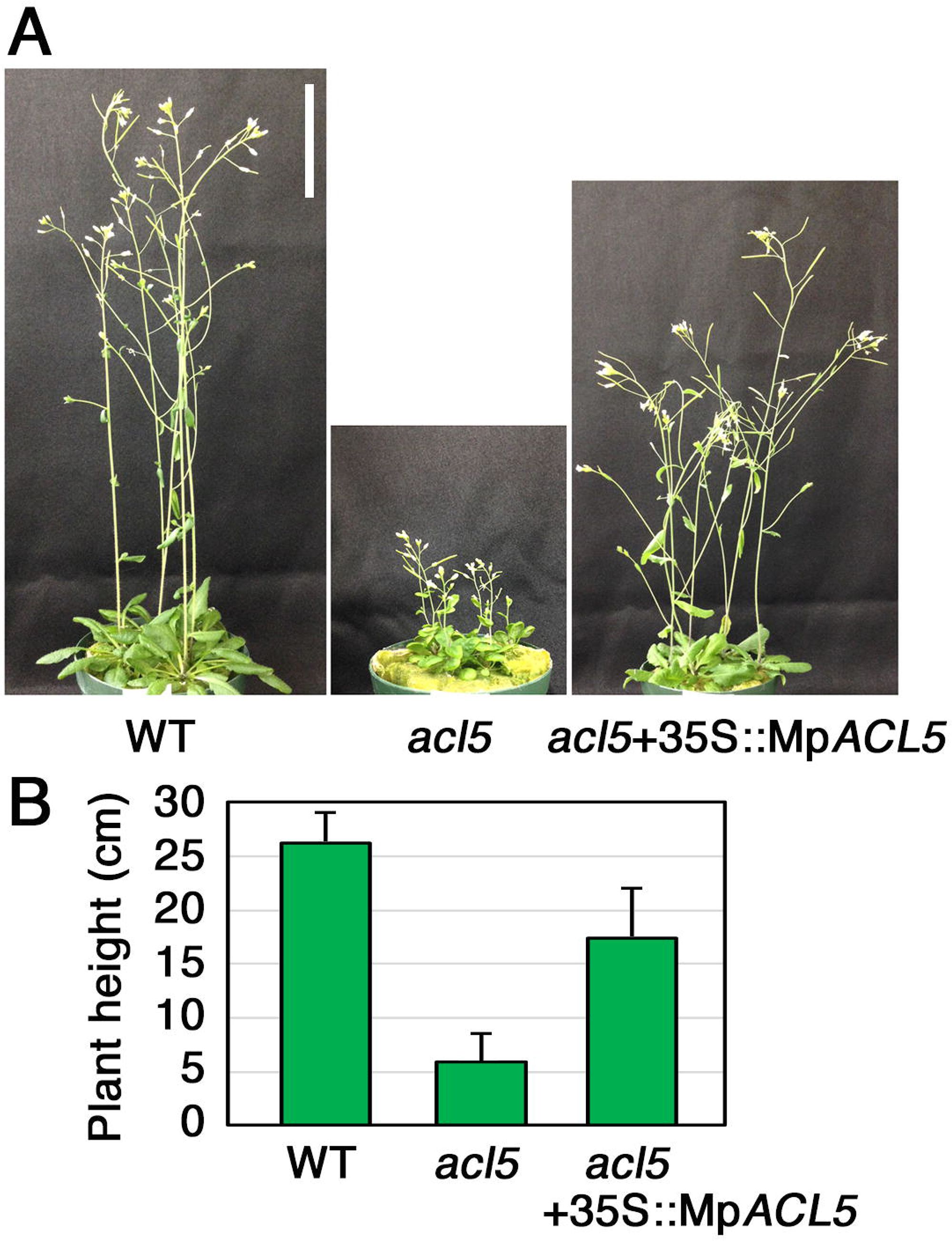
Complementation of the *Arabidopsis acl5* phenotype by Mp*ACL5*. (A) Phenotypes of wild-type Col-0 (WT), *acl5-1*, and a representative transgenic line of *acl5-1* expressing Mp*ACL5* under the CaMV 35S promoter, #1. Plants were grown for 5 weeks on rockwool placed in vermiculite. Bar = 5 cm. (B) Plant height of 30-day-old plants of the wild-type Col-0, *acl5-1*, and *acl5-1* with the 35S:Mp*ACL5*, #1. Bars indicate SD (n=10).

**Fig. 2.**
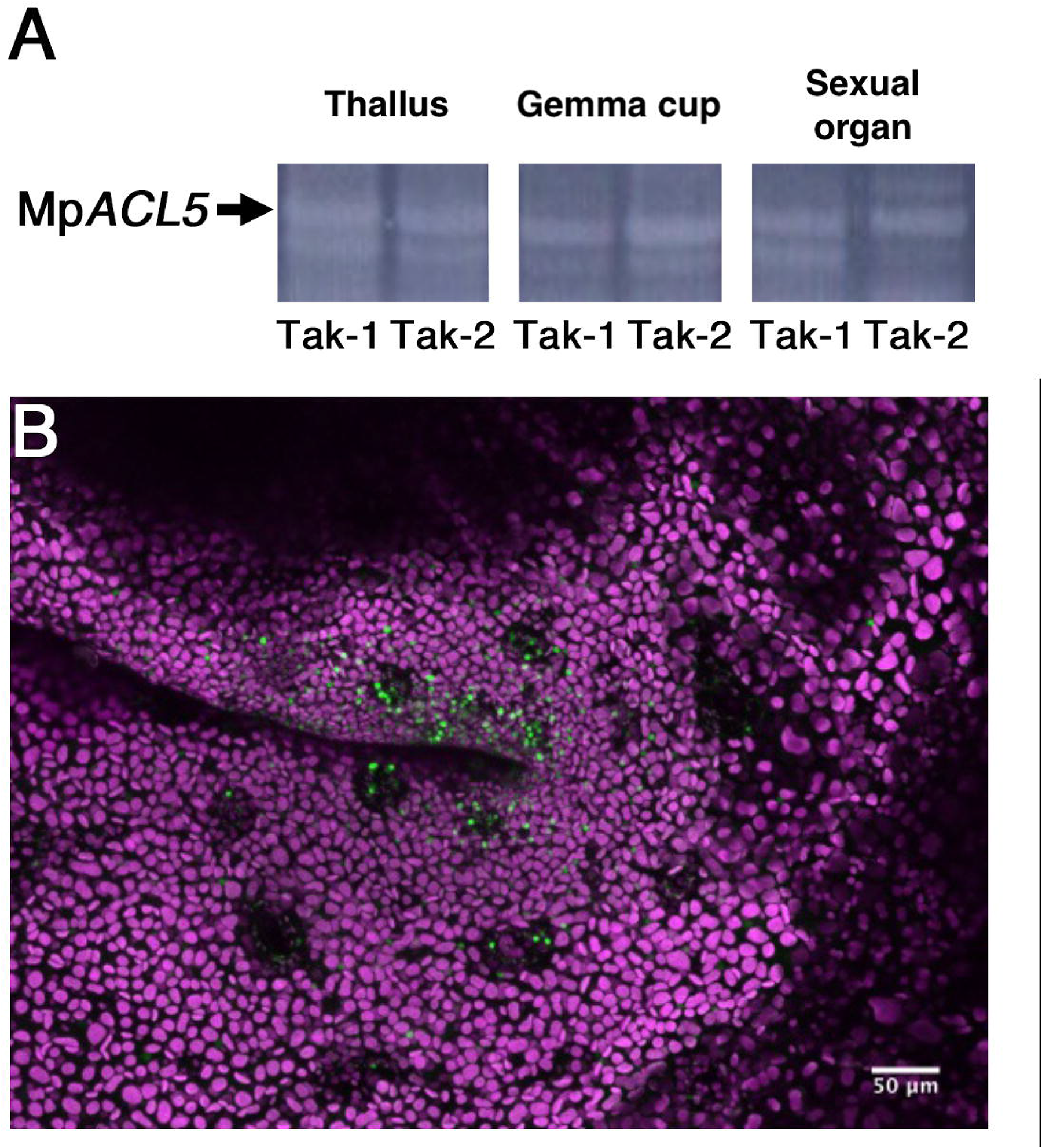
Expression pattern of Mp*ACL5*. (A) RT-PCR analysis of Mp*ACL5* in thallus, gemma cup, and sexual organ. (B) *Citrine-NLS* expression under the Mp*ACL5* promoter. Confocal z-stack image of a notch region indicating the accumulation of Citrine-NLS (green) in the nuclei of thallus cells. Bar = 50 µm.

### Mp*acl5* mutants show abnormal growth

To explore the function of Mp*ACL5*, loss-of-function mutants of Mp*ACL5* were generated by the CRISPR/Cas9 genome editing system (Sugano et al. 2018). The guide RNA was designed from the 95th to 112th bases of the Mp*ACL5* coding sequence (Fig. 3A) that corresponds to the N-terminal region essential for the binding of S-adenosylmethionine and its catalytic activity. We isolated two independent mutant lines, Mp*acl5-1* and Mp*acl5-2*. Mp*acl5-1* harbors a 57-bp deletion, which causes the deletion of 19 amino acid residues including Lys-31 essential for the binding of decarboxylated S-adenosylmethionine and was later found to be a female line. Mp*acl5-2* is a male line with a 19-bp deletion, which results in a frameshift mutation with a premature stop codon and a putative truncated peptide of 102 amino acid residues. Thus, both Mp*acl5-1* and Mp*acl5-2* may represent a null allele.

**Fig. 3.**
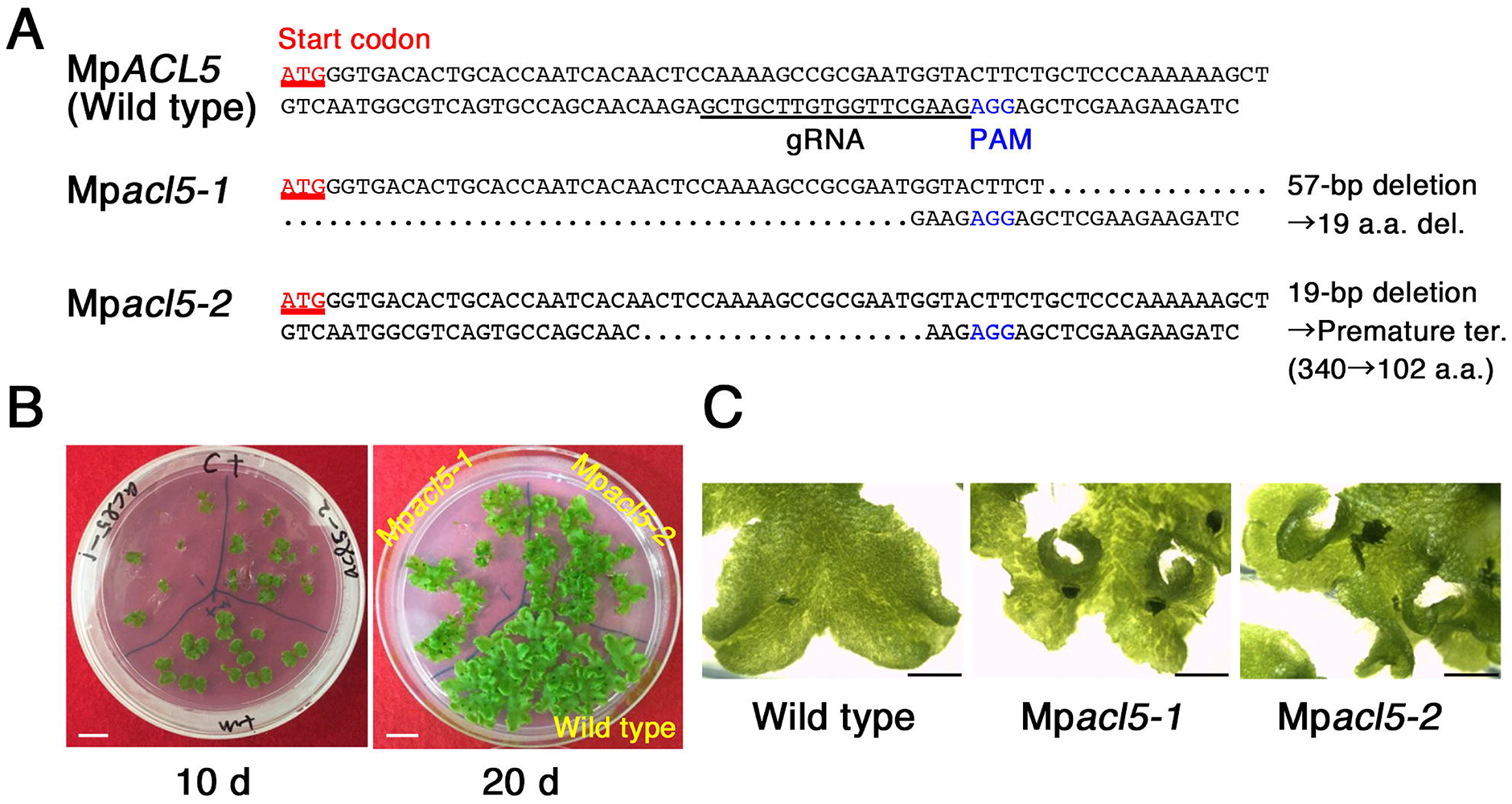
Generation of Mp*acl5* mutants. (A) DNA sequences of Mp*ACL5* around the CAS9 target site. Dots indicate nucleotides deleted by genome editing. (B) Wild-type (Tak-1) and Mp*acl5* mutant thalli grown in the B5 medium for 10 and 20 days. Bars = 1 mm. (C) Magnified views of wild-type (Tak-1) and Mp*acl5* mutant thalli grown in the B5 medium for 20 days. Bars = 1 mm.

Both mutant lines exhibited similar morphological defects in the flat leaf-like vegetative organ, thallus, which grows and periodically bifurcates through the activity of meristematic notch region (Fig. 3B). The thallus has a clear dorsal-ventral morphology: air pores, photosynthetic assimilatory filaments, and gemma cups were formed on the dorsal side whereas rhizoids and ventral scales were generated on the ventral side. Rhizoids elongate by tip growth to form hair-like protrusions. In Mp*acl5* mutants, growth of the thallus was severely retarded and the peripheral region near the meristematic notch was curly and distorted toward the outside of the thallus (Fig. 3C). On the other hand, no morphological abnormalities were observed in gemma cups, air pores, assimilatory filaments, ventral scales, and rhizoids.

Then we successfully induced female and male sexual organs, archegoniophores and antheridiophores, in Mp*acl5-1* and Mp*acl5-2*, respectively, indicating that MpACL5 is not essential for phase transition from vegetative to reproductive growth. However, both mutants showed shorter stalks of gametangiophores than the wild type (Fig. 4A, B). Measurement of the stalk cell length revealed no difference in the cell length between the wild type and Mp*acl5* mutants, suggesting that the suppression of stalk elongation is caused by a decrease in the cell number (Fig. 4C). Furthermore, the mutant stalks exhibited a fasciation phenotype. Wild-type gametangiophores usually develop one by one from a meristematic notch region, whereas the mutant gametangiopores were formed from a notch region in a fusion manner and then branched off (Fig. 4D). Three coalescence patterns were observed depending on where they branch; two gametangiopores were branched at the base of the stalk, in the middle of the stalk, or at the tip of the stalk. Although the fasciation phenotype was observed in both male and female gametangiophores of Mp*acl5*, the fasciation of archegoniophores was more obvious than that of antheridiophores; the former showed three coalescence patterns, whereas the mutant antheridiophores had the stalks branched at the tip. Moreover, while two bundles of pegged rhizoids are formed in a stalk of the wild-type gametangiophore, four bundles of pegged rhizoids were found in both male and female gametangiophores of Mp*acl5* mutants (Fig. 4E), indicating a fusion of stalks. Interestingly, the gametangiophores of Mp*acl5* mutants often initiated secondary thalli from their peripheral tip region and they in turn generated secondary gametangiophores, resulting in the double-decker gametangiophores (Fig. 4F). Under the scanning electron microscopy (SEM), the mutant gametangiophores showed a distorted and rough surface morphology. Digitate rays of archegoniophores were smaller in the mutants than in the wild type. The size reduction and the morphological defects were enhanced in the secondary gametangiophores. The secondary thalli did not form gemma cup. The male secondary thalli generated antheridia on the dorsal side, some of which were exposed from the antheridia pores (Fig. 5), suggesting that the secondary thallus has an intermediate character between vegetative and reproductive organs.

**Fig. 4.**
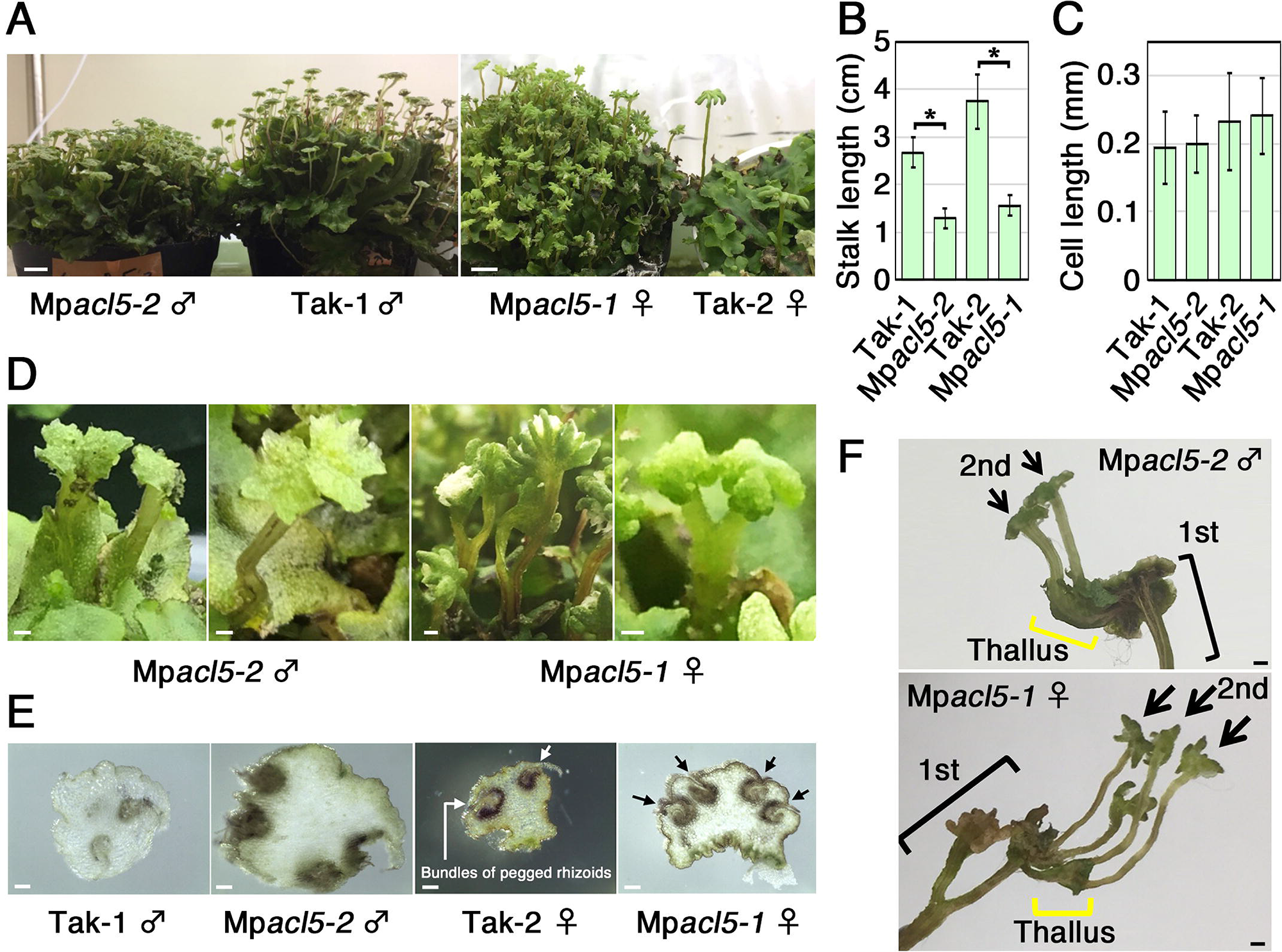
Phenotype of sexual organs in Mp*acl5* mutants. (A) Gross morphology of sexual organs in the wild type and Mp*acl5* mutants. Bars = 1 cm. (B) Length of stalks of sexual organs in the wild type and Mp*acl5* mutants. Bars indicate SD (n=20). Asterisks indicate values determined by Student’s *t* test to be significantly different from the wild type (*P < 0.05). (C) Length of stalk cells of sexual organs in the wild type and Mp*acl5* mutants. Bars indicate SD (n=20). (D) Fasciation phenotype of stalks in Mp*acl5* mutants. Bars = 1 mm. (E) Cross sections of stalks of sexual organs in the wild type and Mp*acl5* mutants. Two bundles of pegged rhizoids (arrows) were observed in the wild type while four bundles were formed in Mp*acl5*. Bars = 100 µm. (F) Secondary sexual organs formed in Mp*acl5*. Both male and female secondary organs (2nd) were generated from the thalli often formed from their respective primary sexual organs (1st). Bars = 1 mm.

**Fig. 5.**
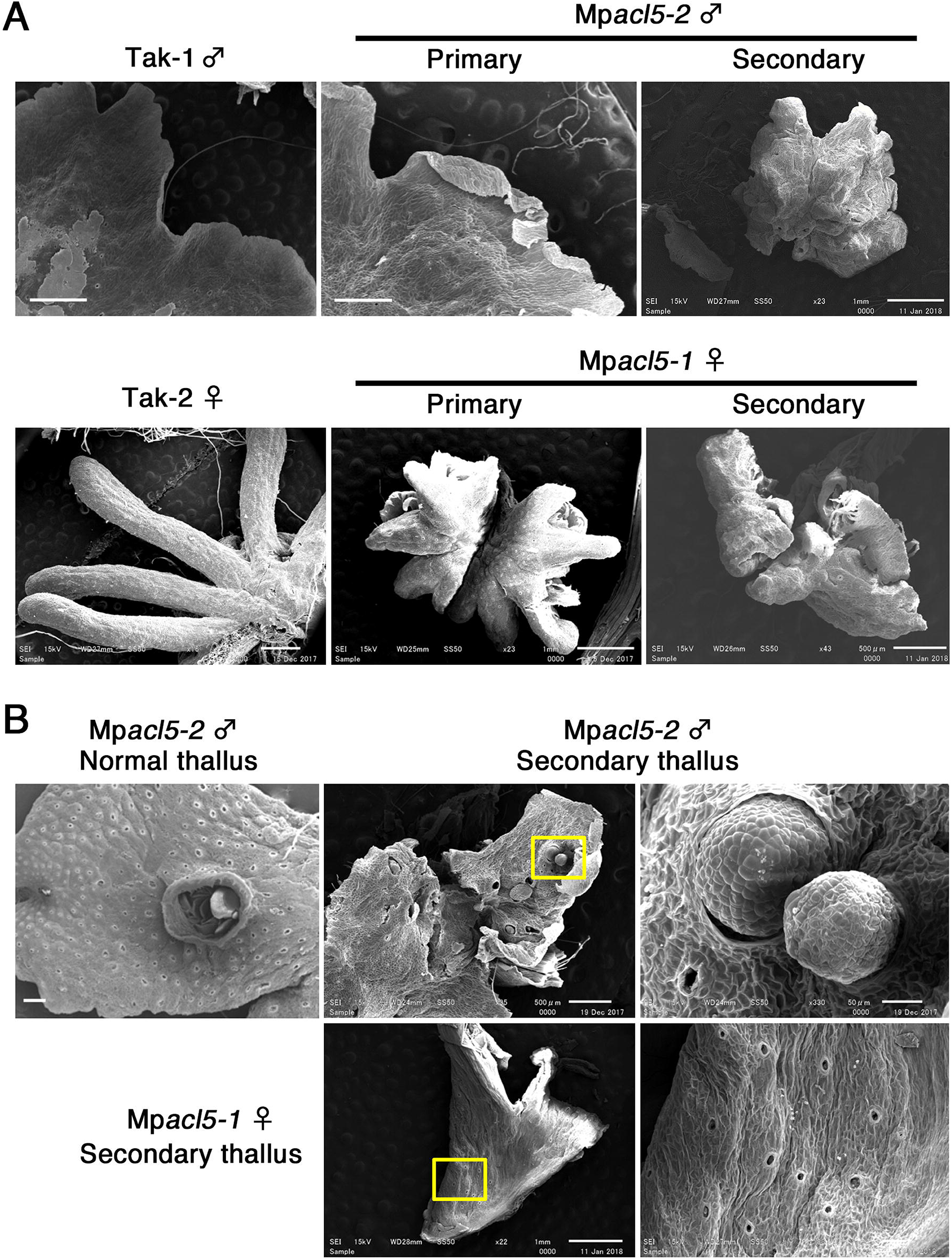
SEM images of gametophores and thalli in the wild type and Mp*acl5* mutants. (A) Morphology of antheridiophores and archegoniophores. Bars = 0.5 mm (upper left, upper center, lower right), 1 mm (upper right, lower left, lower center). (B) Morphology of normal thallus and the secondary thalli generated from the primary sexual organs in Mp*acl5*. Antheridia were formed on the secondary thallus of Mp*acl5*. Bars = 1 mm (upper left, lower center), 0.5 mm (upper center), 50 µm (upper right), 100 µm (lower right).

### Mp*BHLH42* may not be a target of thermospermine

The *M. polymorpha* genome also has a single gene homologous to the *Arabidopsis SAC51*, Mp5g09710 (Mapoly0048s0099, MpBHLH42). Translation of the *SAC51* family is the only known process targeted by thermospermine. Although the 5’ leader region of the Mp*BHLH42* mRNA contains seven AUGs (Supplementary Fig. S3A), none of the three long uORF-encoding peptides are homologous to those conserved among the *SAC51* family in vascular plants (Supplementary Fig. S3B). We further found that the 5’ leader showed no response to thermospermine at least when it was fused to the β-glucuronidase reporter gene and expressed under the CaMV 35S promoter in transgenic *Arabidopsis* plants (Supplementary Fig. S3C). It is thus possible that *M. polymorpha* has different target genes and downstream events of thermospermine from those in vascular plants.

### Mp*acl5* mutants are hypersensitive to stresses

To examine whether exogenous thermospermine can rescue the growth defects of thallus in Mp*acl5*, the thllus was grown in the media supplemented with thermospermine or spermidine, a precursor of thermospermine (Fig. 6). Unexpectedly, both thermospermine and spermidine severely suppressed the growth of thallus in Mp*acl5* but not in the wild type. Spermine was also inhibitory to the growth only in Mp*acl5*. The hypersensitivity of Mp*acl5* mutants to exogenous polyamines suggests a serious effect of thermospermine deficiency on cellular physiology. To examine its effect on gene expression, we performed RNA-seq analysis with the thallus of the wild type and Mp*acl5* and found a large number of genes with altered expression levels in the mutants (Supplementary Table S1). To confirm the RNA-seq data, we subsequently performed quantitative RT-PCR experiments for some genes and found that expression of heat-shock genes, Mp7g06480 (Mapoly0057s0019) and Mp7g07900 (Mapoly0076s0004), and a gene with unknown function, Mp1g09210 (Mapoly0036s0157), was remarkably increased while that of a gene for fucosidase-like protein, Mp5g07800 (Mapoly0127s0004), a gene for Caffeic acid 3-O-methyltransferase-like protein, Mp2g07390 (Mapoly0015s0026, MpOMT8), and a gene with unknown function, Mp8g16710 (Mapoly0030s0004), was down-regulated in Mp*acl5* (Fig. 7).

**Fig. 6.**
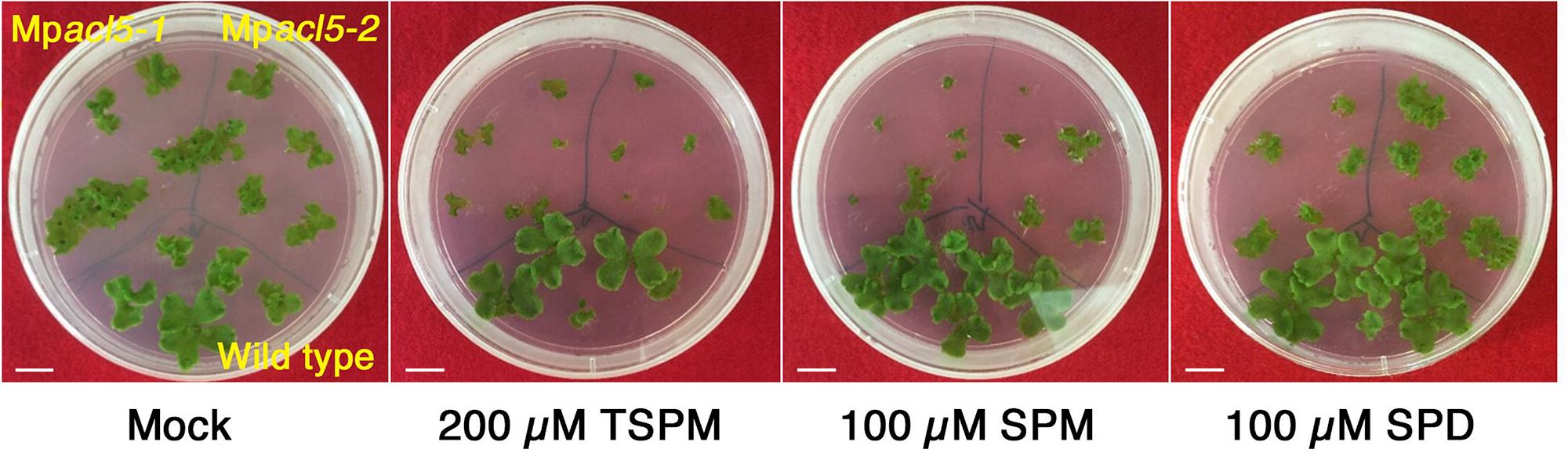
Effect of polyamines on the growth of wild-type (Tak-1) and Mp*acl5* mutant thalli. Gemmae were planted in the B5 medium supplemented with thermospermine (TSPM), spermine (SPM), or spermdine (SPD), and grown for 20 days. Bars = 1 cm.

**Fig. 7.**
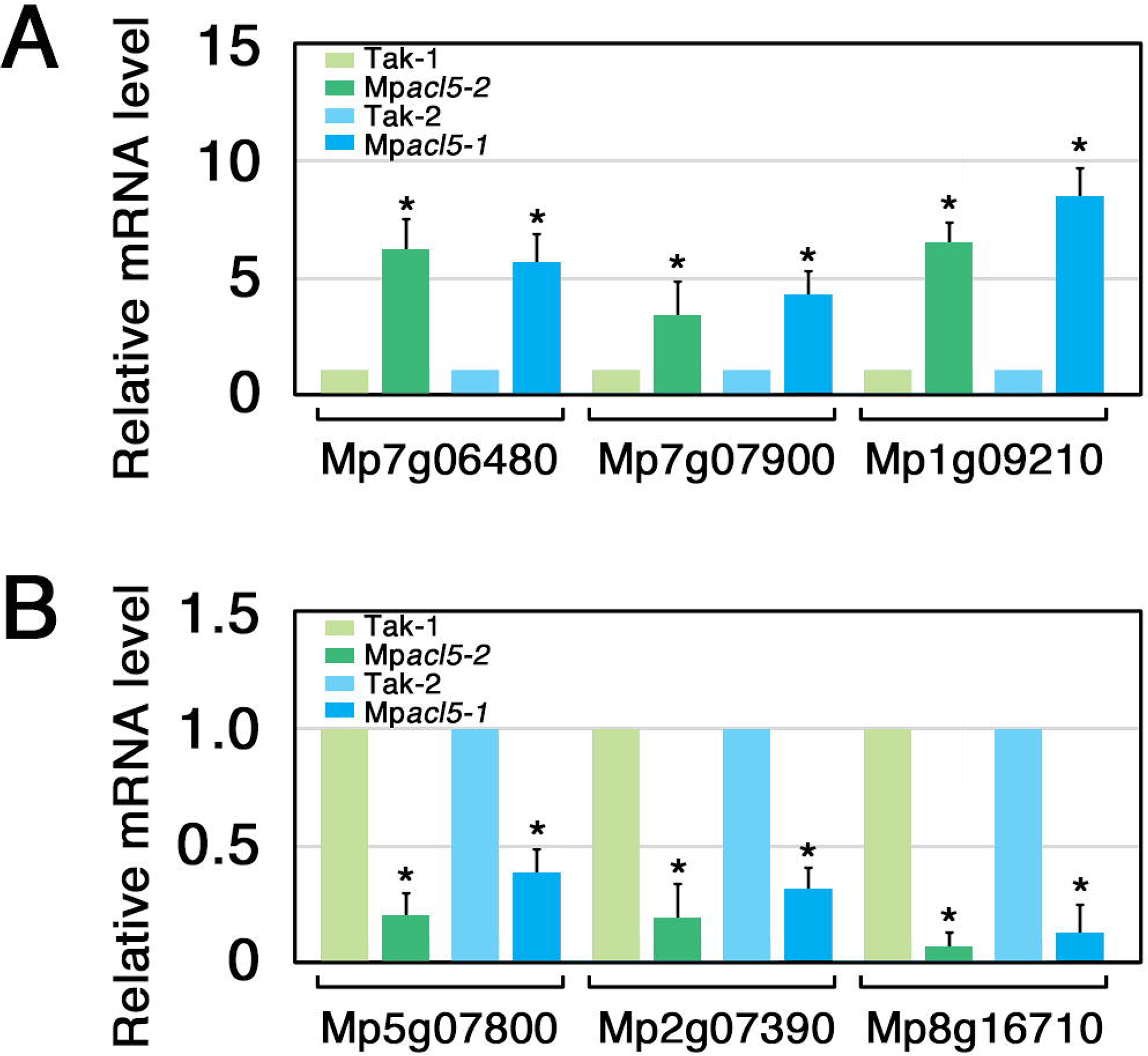
Gene expression altered in the Mp*acl5* mutants. RT-qPCR analysis was conducted to confirm the RNA-seq data, which showed a set of genes upregulated or downregulated in the Mp*acl5* mutant thalli. Relative mRNA levels of representative three genes increased (A) and decreased (B) in Mp*acl5* are shown. Mp7g06480 and Mp7g07900 encode small heat-shock proteins. Mp1g09210 and Mp8g16710 encode a protein with unknown function. Mp5g07800 and Mp2g07390 encode fucosidase-like protein and caffeic acid 3-O-methyltransferase-like protein, respectively. Bars indicate SD (n=3). Asterisks indicate values determined by Student’s *t* test to be significantly different from the wild type (*P < 0.05).

We then analyzed the response to heat stress in Mp*acl5*. The thalli were treated at 37 °C for 6 h and further grown at 22 °C for 5 days. While the wild-type thalli still survived and continued to grow, the mutant thalli showed severe chlorosis and eventually died (Fig. 8A), suggesting that the increased expression of heat-shock genes in Mp*acl5* is not sufficient for the survival of the mutants. These results and the absence of spermine synthase genes in bryophytes raised us a possibility that thermospermine is involved in stress tolerance in *M. polymorpha*. We thus examined the salt sensitivity of Mp*acl5*. When grown on the medium supplemented with 200 mM NaCl, the thalli of the Mp*acl5* mutants showed more severe chlorosis compared to those of the wild type (Fig. 8B, C), suggesting the requirement for thermospermine in salt stress response.

**Fig. 8.**
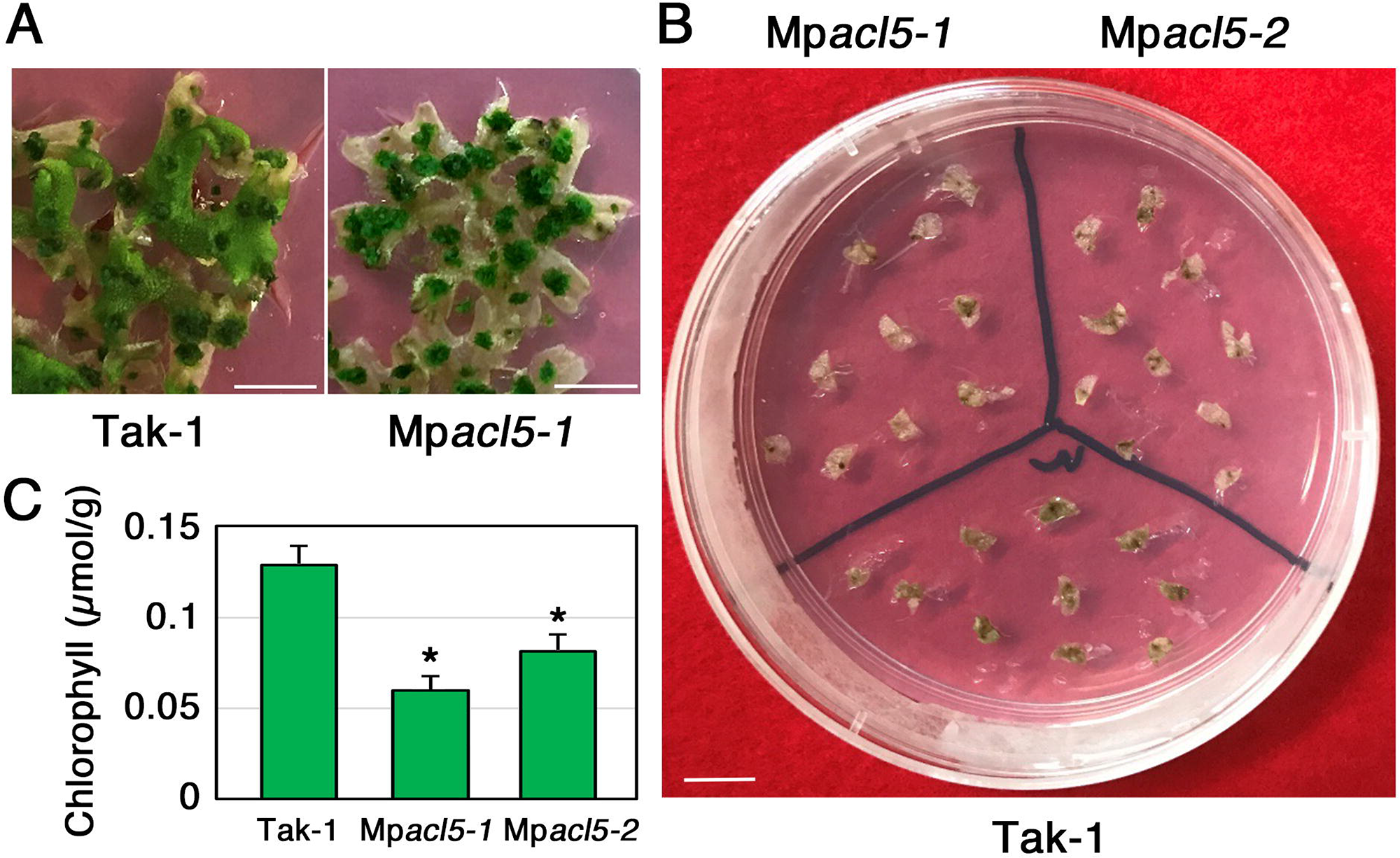
Hypersensitivity of Mp*acl5* mutants to heat and salt stress. (A) Effect of heat stress on the growth of the wild type and Mp*acl5* mutants. Gemmalings of the wild type and Mp*acl5* mutants were grown for 4 weeks on the B5 agar medium in 22°C, treated with 37°C for 6 hours, and grown for 5 days in 22°C. Bars = 1 cm. (B) Effect of NaCl on the growth of the wild type and Mp*acl5* mutants. Gemmalings of the wild type and Mp*acl5* mutants were grown for 7 days on the B5 agar medium without NaCl, transferred to the medium supplemented with 200 mM NaCl, and grown for 12 days. Bar = 1 cm. (C) Chlorophyll content in the wild type and Mp*acl5* mutants grown in the presence of 200 mM NaCl as in (B). Bars indicate SD (n=3). Asterisks indicate values determined by Student’s *t* test to be significantly different from the wild type, Tak-1 (*P < 0.05).

## Discussion

In *Arabidopsis* and probably in other vascular plants, *ACL5* expression is induced by auxin and thermospermine is a key signal for avoiding excessive xylem development in the negative feedback system of auxin-dependent vascular formation. In non-vascular bryophytes, the role of thermospermine remained to be addressed. Here we generated knockout mutants of Mp*ACL5*. Mp*ACL5* is expressed in actively dividing cells but not in specific cell types. In accordance with the expression pattern, Mp*acl5* mutants show growth suppression of vegetative thalli and reproductive gametangiophores rather than defects of tissue and cellular differentiation (Fig. 3). Thus, thermospermine appears to act as a growth-promoting factor in the basal land plant, contrasting to the role as a suppressor of xylem development in angiosperms.

The gametangiophores of Mp*acl5* mutants had short stalks and an increased number of bundles of pegged rhizoids, which are thought to serve as a conductive tissue (Fig. 4). These morphological defects are reminiscent of the *Arabidopsis acl5* mutant which has short inflorescence stems with excess xylems. However, it needs some caution on this morphological similarity because these defects are caused by different mechanisms. The dwarf phenotype of gametangiophores is due to reduced cell numbers rather than reduced cell elongation, which is observed in *Arabidopsis acl5* (Hanzawa et al. 1997). The increased rhizoid bundles are resulted from fasciation of two gametangiophores rather than excess formation of rhizoids (Fig. 4). Thus, it is better to conclude that thermospermine is a major regulator of gametangiophore development to promote stalk elongation and to suppress fasciation of gametangiophores. In the latter function, thermospermine may limit gametangiophore formation for single gametangiophore to generate from one meristematic notch region because of two gametangiophores generated from a notch region in Mp*acl5* mutants. This could be correlated with another function of thermospermine to suppress indeterminate growth of gametangiophores.

According to previous anatomical studies, the gametangiopore retains morphological characters of thallus and may be derived from its morphological transformation. In Mp*acl5* mutants, gametangiopores appear to dedifferentiate into vegetative thalli, which in turn develop secondary gametangiopores (Fig. 5). In summary, thermospermine may have opposite functions to promote growth of thallus and stalks and to suppress gametangiopore formation and dedifferentiation of gametangiopore into thallus. The molecular mechanisms of these functions remain obscure but are likely to be mediated by translational or transcriptional controls as suggested from our present RNA-seq analysis and previous studies in *Arabidopsis* (Kakehi et al. 2008, Cai et al. 2016, Ishitsuka et al. 2019).

Recent studies indicate that not only spermine but also thermospermine is involved in stress tolerance in angiosperms (Sagor et al. 2012, Marina et al. 2013, Mo et al. 2015, Zarza et al. 2017). *Arabidopsis acl5* mutant shows high salt sensitivity caused by increase of xylem formation and salt uptake (Shinohara et al. 2019). In agreement with these studies, Mp*acl5* mutants are also susceptible to salt stress (Fig. 8). Generally, gametophytes in bryophytes have been thought to absorb water and nutrients from their surface rather than specific conductive tissues. Thus, the hypersensitivity to salt stress could be attributed to the reduced tolerance to salt stress of Mp*acl5* mutants rather than the increased uptake of salt. This is consistent with their hypersensitive phenotype to heat stress. Taking the increased expression of *HSP* genes into account, the deficiency of thermospermine may constitutively stimulate stress response pathways in Mp*acl5* mutants. This possibility does not exclude direct functions of thermospermine in conferring tolerance to harsh environments as proposed in thermophilic bacteria containing thermospermine and various uncommon polyamines (Sakamoto et al. 2022).

Our present genetic analysis of Mp*ACL5* revealed fundamental function of thermospermine in both organ development and stress tolerances in the basal land plant. Bryophytes do not have spermine and spermine synthase (SPMS) gene. Phylogenetic analyses imply that SPMS might be acquired from spermatophytes during land plant evolution (Minguet et al. 2008). In this scenario, it is possible that spermine takes over the original function of thermospermine in stress responses and then thermospermine is coopted into specific regulation of development and differentiation of cells and tissues. Thus, our study implies the evolutionarily ancient multiple functions of thermospermine required for adaptation to the terrestrial environment during land plant evolution. Further investigation in algae and other basal land plants will provide insight into the functional origin and diversity of this primordial phytohormone.

## Materials and Methods

### Plant Material and Growth Conditions

*Marchantia polymorpha* accessions Takaragaike-1 (Tak-1, male) and Takaragaike-2 (Tak-2, female) were used as the wild type. Thallus, gemmalings, and sporelings were grown on the half-strength Gamborg’s B5 medium solidified with 1% agar at 22°C under continuous white light. Gametangiophore was induced under continuous white light with far-red irradiation (Chiyoda et al., 2008). For polyamine treatment, spermidine, spermine, and thermospermine (Santa Cruz Biotechnology, TX, USA) were dissolved in water and added to the B5 medium. Dormant gemmae were planted on the B5 medium supplemented with each polyamine.

For *Arabidopsis* experiments, the Columbia (Col-0) accession was used as the wild type. Plants were grown on rockwool as described previously (Hanzawa et al. 1997).

### Complementation of *Arabidopsis acl5-1*

The full-length MpACL5 amplified from Tak-1 cDNA by PCR using TaKaRA EX Taq (Takara, Shiga, Japan) with the primers, FXb (TCTAG AATGG GTGAC ACTGC ACCA) and RBm (GGATC CTAAT GGGAT TTCGC ATTGG), was cloned into pGEM-T Easy cloning vector (Promega, WI, USA). The fragment was then excised by *Xba*I and *Bam*HI and inserted into the downstream of the CaMV35S promoter of pBI121 (Takara) whose GUS reporter gene was removed in advance to generate the CaMV35S:MpACL5 construct. The resulting Ti plasmid was introduced into *Agrobacterium* GV3101 (pMP90) strain by electroporation and the T-DNA was introduced into the *Arabidopsis acl5-1* mutant in the Col-0 background (Hanzawa et al. 1997) by the floral dip method (Clough and Bent 1998). Transformants were selected with 40 µg/mL kanamycin and 100 µg/mL cefotaxime and T3 lines were analyzed in further experiments.

### Promoter activity of Mp*ACL5*

To construct Mp*ACL5pro:Citrine-NLS*, MpACL5 genomic region, including a 3324-b region upstream of the initiation codon and 24-b coding region corresponding to 8-amino-acid length, was amplified from Tak-1 genomic DNA by PCR using KOD plus (Toyobo, Osaka, Japan) with the primers, MpACL5pro-F-3k (CACC GAT ACA CGG CTC ATG TTG AAA ATT AG) and MpACL5-R-8aa (TGT GAT TGG TGC AGT GTC ACC CAT), and cloned into pENTR/D-TOPO (Thermo Fisher Scientific, MA, USA). This entry vector was used in the LR reaction with the Gateway binary vector pMpGWB115 (Ishizaki et al. 2015) to generate the Mp*ACL5pro:Citrine-NLS* construct, in which nuclear localized Citrine was translationally fused with the first 8-amino-acid sequence of MpACL5. This vector was introduced into *Agrobacterium* GV3101 (pMP90) strain by electroporation and introduced into regenerating thalli of Tak-1 as previously described (Kubota et al. 2013). Transformants were selected with 10 µg/mL hygromycin B and 100 µg/mL cefotaxime.

### Bacterial MpACL5 production

The above-described cloned full length MpACL5 fragment was excised by *Xba*I and *Hind*III and transferred into the pMal-c2 vector (New England BioLabs, MA, USA) to generate pMal-MpACL5 construct. pMal-MpACL5, pMal-AtACL5 (Hanzawa et al. 2000), and empty vector were introduced into *E. coli* DH5α. The transformed *E. coli* was cultured in 2 mL LB medium supplemented with 2% glucose and 100 µg/mL ampicillin for 6 h and then cultured in the presence of 100 µM IPTG for 3 h to induce maltose binding protein-ACL5 fusion proteins. After centrifugation at 15,000 rpm for 1 min, precipitate was suspended with 800 µL of 5% PCA, sonicated by a SONIFIER 250 (EMERSON BRANSON, CT, USA) for 1 min in the constant mode output 2 on ice, and centrifuged at 15,000 rpm at 4°C. 400 µL of supernatant was neutralized with 200 µL 2N NaOH and treated with 2 µL benzoyl chloride at room temperature for 20 min. 400 µL saturated sodium chloride was then added, followed by addition of 400 µL diethyl ether and vigorous mixing. After centrifugation at 4°C 3000 g for 10 min, the organic layer was collected in 2 mL microtubes, evaporated, and resuspended in 50 µL of methanol to make a polyamine extract. Polyamines were detected by using a reverse phase HPLC system equipped with TSKgel ODS-80Ts column (Toso, Tokyo, Japan)(Tong et al. 2014).

### RT-qPCR

Total RNA was isolated from the thallus, gemma cup, and gametandiopore of one-month-old Tak-1 and Tak-2 by NucleoSpin RNA Plant (Takara) or Monarch Total RNA Miniprep Kit (New England BioLabs) according to the manufactures’ instruction. Thalli and gemma cup separated from the thalli by a scalpel were obtained from 20-day-old wild type plants. Sexual organs were obtained from one-month-old wild type plants. These were immediately frozen in liquid N_2_ for subsequent RNA extraction. For each sample, 0.5 µg of total RNA was reverse transcribed to cDNA using ReverTra Ace reverse transcriptase (Toyobo) according to the manufacture’s protocol. PCR was performed by TaKaRa EX Taq (Takara) and primers, Mp*ACL5* ex2.5 (AAGCT ACAGT CAGCT GAG) and ex4R (CAAGA GTCGG CGTAC GAT) in the thermal cycle of 95°C 2 min – 40 cycles of 95L 30 sec, 55L 30 sec, 72L 90 sec. Real-time PCR was performed on a thermal cycler Dice Real Time System (Takara) using KAPA SYBR FAST qPCR Kit (KAPA Biosystems) according to the manufacturer’s method. Transcript levels of Mp*EF1α* or Mp*ACT7* were used as a reference for normalization (Kubota et al. 2014, Saint-Marcoux et al. 2015). Primers used in RT-qPCR are listed in Supplementary Table S2.

### RNA-seq

RNA-seq was conducted according to Yamaoka et al. (2018). Total RNA was isolated from 14-day-old thalli of the wild type and Mp*acl5* mutants grown on the half-strength B5 agar medium using NucleoSpin RNA Plant (Takara). The sequence libraries were generated by TruSeq RNA Sample Prep Kit (Illumina, CA, USA) and sequenced in Illumina HiSeq 1500 platform. The mapping of sequence reads and gene expression analysis were also conducted according to Yamaoka et al. (2018). Four biological replicates were used for RNA-seq analysis.

### Mutagenesis of Mp*ACL5*

For the CRISPR/Cas9 construct of MpACL5, complementary DNA oligos encoding 18-base target sequences of gRNA, gRNA-F (CTCG GCT GCT TGT GGT TCG AAG) and gRNA-R (AAAC CTT CGA ACC ACA AGC AGC), were annealed and cloned into the *Bsa*I site of an entry vector pMpEn_03 (Sugano et al. 2018). The MpU6-1pro:gRNA in pMpEn_03 was transferred into a binary vector pMpGE010 by LR reaction using Gateway LR Clonase II enzyme mix (Thermo Fisher Scientific). The resulting binary vector was introduced into *Agrobacterium* GV3101 (pMP90) strain by electroporation. The construct was introduced into wild-type sporelings (F1 spores produced by crossing Tak-2 and Tak-1) by *Agrobacterium*-mediated transformation method (Ishizaki et al. 2008). Plants were transferred on the half B5 agar medium containing 10 µg/mL hygromycin and 100 µg/mL cefotaxime. For the selection of mutants, genomic DNA of transformants was extracted and subjected to PCR by using a DNA polymerase KOD FX neo (Toyobo) and gene-specific primers to amplify DNA flanking the target sequence. The primers used for PCR are shown in Supplementary Table S2. By using the amplified DNA as a template, the sequencing reaction was performed by BigDye terminator ver.3.1 (Thermo Fisher Scientific) and a gene-specific primer (Supplementary Table S2). DNA sequence was analyzed by ABI3500 Genetic Analyzer (Applied Biosystems).

### Microscopy

Morphology of plants was observed by using a stereoscopic microscope S8APO0 equipped with a CCD camera DFC500 or a light microscope DM5000B equipped with DFC500 (Leica Microsystems, Wetzlar, Germany). The whole morphology of plants and gametangiophores were photographed by a single-lens reflex camera D5600 (Nikon, Tokyo Japan).

The thalli and reproductive organs were fixed overnight in a 1% aldehyde solution (glutaraldehyde 1%, phosphate buffer 5%), and substituted by ethanol while gradually increasing the ethanol concentration from 50% to 100%. 100% ethanol was replaced with isoamyl acetate, dried in a critical point dryer JCPD-5 (JEOL, Tokyo, Japan). After drying, the samples were coated by gold evaporation with JFC-1200 Fine Coater (JEOL) and observed under a scanning electron microscope JSM-6510LV (JEOL).

To analyze promoter activity of Mp*ACL5*, Mp*ACL5pro:Citrine-NLS* plants were grown on half-strength B5 medium for 5 to 7 days under continuous white light. The gemmalings were placed on a glass slide with small aliquots of water, covered with a glass strip, and observed under a FV1200 confocal laser-scanning microscope (Evident, Tokyo, Japan) equipped with a high-sensitivity GaAsP detector and silicone oil objective lenses [Evident, 30 x, numerical aperture (NA) = 1.05; 60 x, NA = 1.3,]. Silicone oil (SIL300CS, Evident) was used as immersion medium for objective lens. The samples were excited at 473 nm and 559 nm (laser diode). The emission was separated using a FV12-MHSY SDM560 filter (490-540 nm, 575-675 nm, Evident). The images were analyzed by ImageJ (National Institutes of Health).

### Measurement of chlorophyll

Chlorophyll was extracted from 50 mg of thalli in 1 mL of *N,N*-dimethyl formamide at 4°C overnight in the dark and assayed as described (Porra et al. 1989).

## Acknowledgements

We thank Dr. Kimitsune Ishizaki (Kobe University) for useful advice on the transformation of *M. polymorpha* and early genome information and Dr. Masaki Shimamura (Hiroshima Univ.) for discussion and advice on the morphology of *M. polymorpha*.

## Author contributions

T. F., H. M. and T. T. designed the research. T. F., K. I., S. Y., and H. M. performed experiments. T. F., K. I., S. Y., T. K., H. M. and T. T. analyzed data and wrote the paper. H. M. and T. T. supervised the research.

## References

Althoff, F., Kopischke, S., Zobell, O., Ide, K., Ishizaki, K., Kohchi, T. and Zachgo, S. (2014) Comparison of the MpEF1α and CaMV35 promoters for application in Marchantia polymorpha overexpression studies. Transgenic Res. 23: 235–244.

Bowman, J.L. (2016) A brief history of *Marchantia* from greece to genomics. Plant Cell Physiol. 57: 210–229.

Bowman, J.L., Kohchi, T., Yamato, K.T., Jenkins, J., Shu, S., Ishizaki, K., Yamaoka, S., Nishihama, R., Nakamura, Y., Berger, F., **et al**. (2017) Insights into land plant evolution garnered from the *Marchantia polymorpha* genome. Cell 171: 287–304.

Cai, Q., Fukushima, H., Yamamoto, M., Ishii, N., Sakamoto, T., Kurata, T., Motose, H. and Takahashi, T. (2016) The *SAC51* family plays a central role in thermospermine responses in *Arabidopsi*s. Plant Cell Physiol. 57: 1583–1592.

Chiyoda, S., Ishizaki, K., Kataoka, H., Yamato, K.T. and Kohchi, T. (2008) Direct transformation of the liverwort *Marchantia polymorpha* L. by particle bombardment using immature thalli developing from spores. Plant Cell Rep. 27: 1467–1473.

Clay, N.K. and Nelson, T. (2005) Arabidopsis *thickvein* mutation affects vein thickness and organ vascularization, and resides in a provascular cell-specific spermine synthase involved in vein definition and in polar auxin transport. Plant Physiol. 138: 767–777.

Clough, S.J. and Bent, A.F. (1998) Floral dip: a simplified method for Agrobacterium-mediated transformation of *Arabidopsis thaliana*. Plant J. 16: 735–743.

Flores-Sandoval, E., Eklund, D.M. and Bowman, J.L. (2015) A simple auxin transcriptional response system regulates multiple morphogenetic processes in the Liverwort *Marchantia polymorpha*. PLoS Genet. 28: e1005207.

Hanzawa, Y., Takahashi, T. and Komeda, Y. (1997) *ACL5*: an Arabidopsis gene required for internodal elongation after flowering. Plant J. 12: 863–874.

Hanzawa, Y., Takahashi, T., Michael, A.J., Burtin, D., Long, D., Pineiro, M., Coupland, G., Komeda, Y. (2000) *ACAULIS5*, an *Arabidopsis* gene required for stem elongation, encodes a spermine synthase. EMBO J. 19: 4248–4256.

Igarashi, K. and Kashiwagi, K. (2010) Modulation of cellular function by polyamines. Int. J. Biochem. Cell Biol. 42: 39–51.

Imai, A., Akiyama, T., Kato, T., Sato, S., Tabata, S., Yamamoto, K.T. and Takahashi, T. (2004) Spermine is not essential for survival of *Arabidopsis*. FEBS Lett. 556: 148–152.

Imai, A., Hanzawa, Y., Komura, M., Yamamoto, K.T., Komeda, Y. and Takahashi, T. (2006) The dwarf phenotype of the *Arabidopsis acl5-1* mutant is suppressed by a mutation in an upstream ORF of a bHLH gene. Development 133: 3575–3585.

Ishitsuka, S., Yamamoto, M., Miyamoto, M., Kuwashiro, Y., Imai, A., Motose, H. and Takahashi, T. (2019) Complexity and conservation of thermospermine-responsive uORFs of *SAC51* family genes inangiosperms. Front. Plant Sci. 10: 564.

Ishizaki, K., Chiyoda, S., Yamato, K.T. and Kohchi, T. (2008) Agrobacterium-mediated transformation of the haploid liverwort *Marchantia polymorpha* L., an emerging model for plant biology. Plant Cell Physiol. 49: 1084–1091.

Ishizaki, K., Nonomura, M., Kato, H., Yamato, K.T. and Kohchi, T. (2012) Visualization of auxin-mediated transcriptional activation using a common auxin-responsive reporter system in the liverwort *Marchantia polymorpha*. J. Plant Res. 125: 643–651.

Ishizaki, K., Mizutani, M., Shimamura, M., Masuda, A., Nishihama, R. and Kohchi, T. (2013) Essential role of the E3 ubiqutin ligase nopperabo1 in schizogenous intercellular space formation in the liverwort *Marchantia polymorpha*. Plant Cell 25: 4075–4084.

Ishizaki, K., Nishihama, R., Ueda, M., Inoue, K., Ishida, S., Nishimura, Y., Shikanai, T. and Kohchi, T. (2015). Development of gateway binary vector series with four different selection markers for the liverwort *Marchantia polymorpha*. PLoS One 10, e0138876.

Kakehi, J.I., Kuwashiro, Y., Niitsu, M. and Takahashi, T. (2008) Thermospermine is required for stem elongation in *Arabidopsis thaliana*. Plant Cell Physiol. 49: 1342–1349.

Katayama, H., Iwamoto, K., Kariya, Y., Asakawa, T., Kan, T., Fukuda, H. and Ohashi-Ito, **K.** (2015) A negative feedback loop controlling bHLH complexes is involved in vascular cell division and differentiation in the root apical meristem. Curr. Biol. 25: 3144–3150.

Kato, H., Ishizaki, K., Kouno, M., Shirakawa, M., Bowman, J.L., Nishihama, R. and Kohchi, T. (2015) Auxin-mediated transcriptional system with a minimal set of components is critical for morphogenesis through the life cycle in *Marchantia polymorpha*. PLoS Genet. 28: e1005084.

Kubota, A., Ishizaki, K., Hosaka, M. and Kohchi, T. (2013) Efficient *Agrobacterium*-mediated transformation of the liverwort *Marchantia polymorpha* using regenerating thalli. Biosci. Biotechnol. Biochem. 77: 167–172.

Kubota, A., Kita, S., Ishizaki, K., Nishihama, R., Yamato, K.T. and Kohchi, T. (2014) Co-option of a photoperiodic growth-phase transition system during land plant evolution. Nat. Commun. 5: 3668.

Kumar, S., Stecher, G. and Tamura, K. (2016) MEGA7: Molecular Evolutionary Genetics Analysis version 7.0 for bigger datasets. Mol. Biol. Evol. 33: 1870–1874.

Marina, M., Sirera, F.V., Rambla, J.L., Gonzalez, M.E., Blázquez, M.A., Carbonell, J. Pieckenstain, F.L. and Ruiz, O.A. (2013) Thermospermine catabolism increases *Arabidopsis thaliana* resistance to *Pseudomonas viridiflava*. J. Exp. Bot. 64: 1393–1402.

Minguet, E.G., Vera-Sirera, F., Marina, A., Carbonell, J. and Blázquez, M.A. (2008) Evolutionary diversification in polyamine biosynthesis. Mol. Biol. Evol. 25: 2119–2128.

Mo, H., Wang, X., Zhang, Y., Yang, J. and Ma, Z. (2015) Cotton *ACAULIS5* is involved in stem elongation and the plant defense response to *Verticillium dahliae* through thermospermine alteration. Plant Cell Rep. 34: 1975–1985.

Oshima, T. (1979) A new polyamine, thermospermine, 1, 12-diamino-4, 8-diazadodecane, from an extreme thermophile. J. Biol. Chem. 254: 8720–8722.

Pegg, A.E. (2016) Functions of polyamines in mammals. J. Biol. Chem. 291: 14904–14912.

Porra, R.J., Thompson, W.A. and Kriedemann, P.E. (1989) Determination of accurate extinction coefficients and simultaneous equations for assaying chlorophylls a and b extracted with four different solvents: verification of the concentration of chlorophyll II standards by atomic absorption spectroscopy. Biochim. Biophys. Acta 975: 384–394.

Sagor, G.H., Takahashi, H., Niitsu, M., Takahashi, Y., Berberich, T. and Kusano, T. (2012) Exogenous thermospermine has an activity to induce a subset of the defense genes and restrict cucumber mosaic virus multiplication in *Arabidopsis thaliana*. Plant Cell Rep. 31: 1227–1232.

Sakamoto, A., Tamakoshi, A., Moriya, T., Oshima, T., Takao, K., Sugita, Y., Furuchi, T., Niitsu, M., Uemura, T., Igarashi, K. **et al**. (2022) Polyamines produced by an extreme thermophile are essential for cell growth at high temperature. J. Biochem.172: 109–115.

Shimamura, M. (2016) *Marchantia polymorpha*; taxonomy, phylogeny, and morphology of a model system. Plant Cell Physiol. 57: 230–256.

Shinohara, S., Okamoto, T., Motose, H. and Takahashi, T. (2019) Salt hypersensitivity is associated with excessive xylem development in a thermospermine-deficient mutant of *Arabidopsis thaliana*. Plant J. 100: 374–383.

Solé-Gil, A., Hernández-García, J., López-Gresa, M.P., Blázquez, M.A. and Agustí, J. (2019) Conservation of thermospermine synthase activity in vascular and non-vascular plants. Front. Plant Sci. 10: 663.

Sugano, S.S., Nishihama, R., Shirakawa, M., Takagi, J., Matsuda, Y., Ishida, S., Shimada T, Hara-Nishimura, I., Osakabe, K. and Kohchi, T. (2018) Efficient CRISPR/Cas9-based genome editing and its application to conditional genetic analysis in *Marchantia polymorpha*. PLoS One 13: e0205117.

Takahashi, T. and Kakehi, J.I. (2010) Polyamines: ubiquitous polycations with unique roles in growth and stress responses. Ann. Bot. 105: 1–6.

Takahashi, T., Naito, S. and Komeda, Y. (1992) Isolation and analysis of the expression of two genes for the 81-kilodalton heat-shock proteins from *Arabidopsis*. Plant Physiol. 99: 383–390.

Takano, A., Kakehi, J.I. and Takahashi, T. (2012) Thermospermine is not a minor polyamine in the plant kingdom. Plant Cell Physiol. 53: 606–616.

Tiburcio, A.F., Altabella, T., Bitrián, M. and Alcázar, R. (2014) The roles of polyamines during the lifespan of plants: from development to stress. Planta 240: 1–18.

Tong, W., Yoshimoto, K., Kakehi, J.I., Motose, H., Niitsu, M. and Takahashi, T. (2014) Thermospermine modulates expression of auxin-related genes in *Arabidopsis*. Front. Plant Sci. 5: 94.

Vera-Sirera, F., De Rybel, B., Úrbez, C., Kouklas, E., Pesquera, M., Álvarez-Mahecha, J.C., Minguet, E.G., Tuominen, H., Carbonell, J., Borst, J.W. et al. (2015) A bHLH-based feedback loop restricts vascular cell proliferation in plants. Dev. Cell 35: 432–443.

Yamamoto, M. and Takahashi, T. (2017) Thermospermine enhances translation of *SAC51* and *SACL1* in Arabidopsis. Plant Signal. Behav. 12: e1276685. 016.1276685.

Yamaoka, S., Nishihama, R., Yoshitake, Y., Ishida, S., Inoue, K., Saito, M., Okahashi, K., Bao, H., Nishida, H., Yamaguchi, K. **et al**. (2018) Generative cell specification requires transcription factors evolutionarily conserved in land plants. Curr. Biol. 28: 479–486.

Zarza, X., Atanasov, K.E., Marco, F., Arbona, V., Carrasco, P., Kopka, J., Fotopoulos, V., Munnik, T., Gómez-Cadenas, A., Tiburcio, A.F. **et al**. (2017) *Polyamine oxidase 5* loss-of-function mutations in *Arabidopsis thaliana* trigger metabolic and transcriptional reprogramming and promote salt stress tolerance. Plant Cell Environm 40: 527–542.

