## Supplementary Fig. 1-3 for "Thermospermine is an evolutionarily ancestral phytohormone required for organ development and stress responses in the basal land plant *Marchantia polymorpha*"

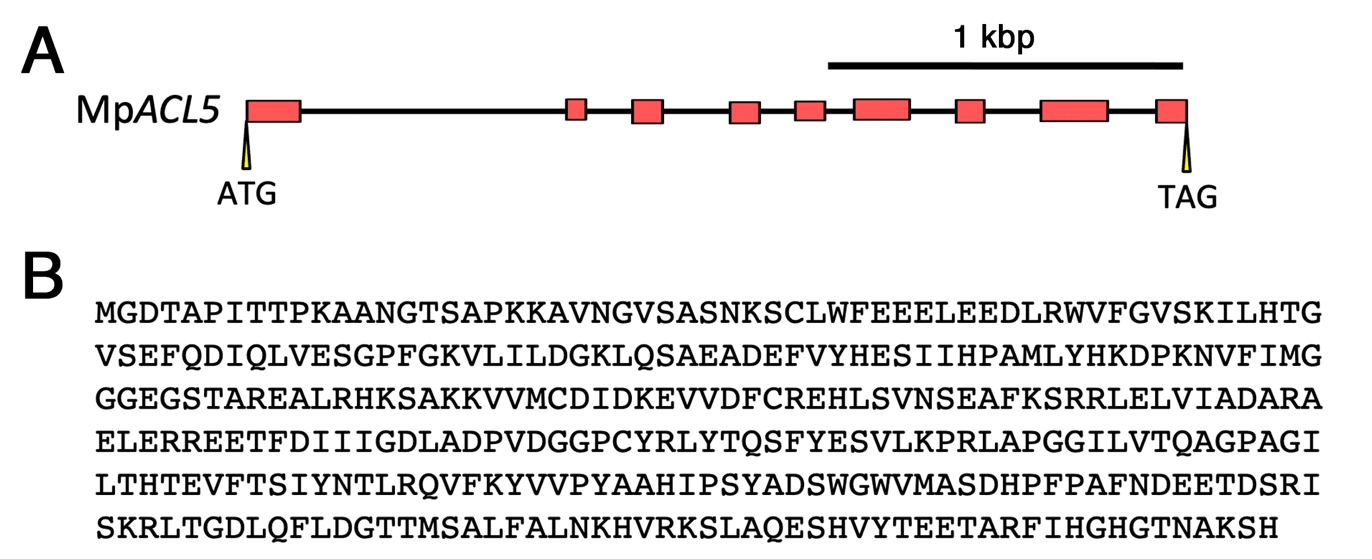

**Supplementary Fig. S1**	Structure of Mp*ACL5*. (A) Exon-intron structure of Mp*ACL5*. Protein coding regions are shown in red boxes. Bars indicate introns. (B) Deduced amino acid sequence of MpACL5.

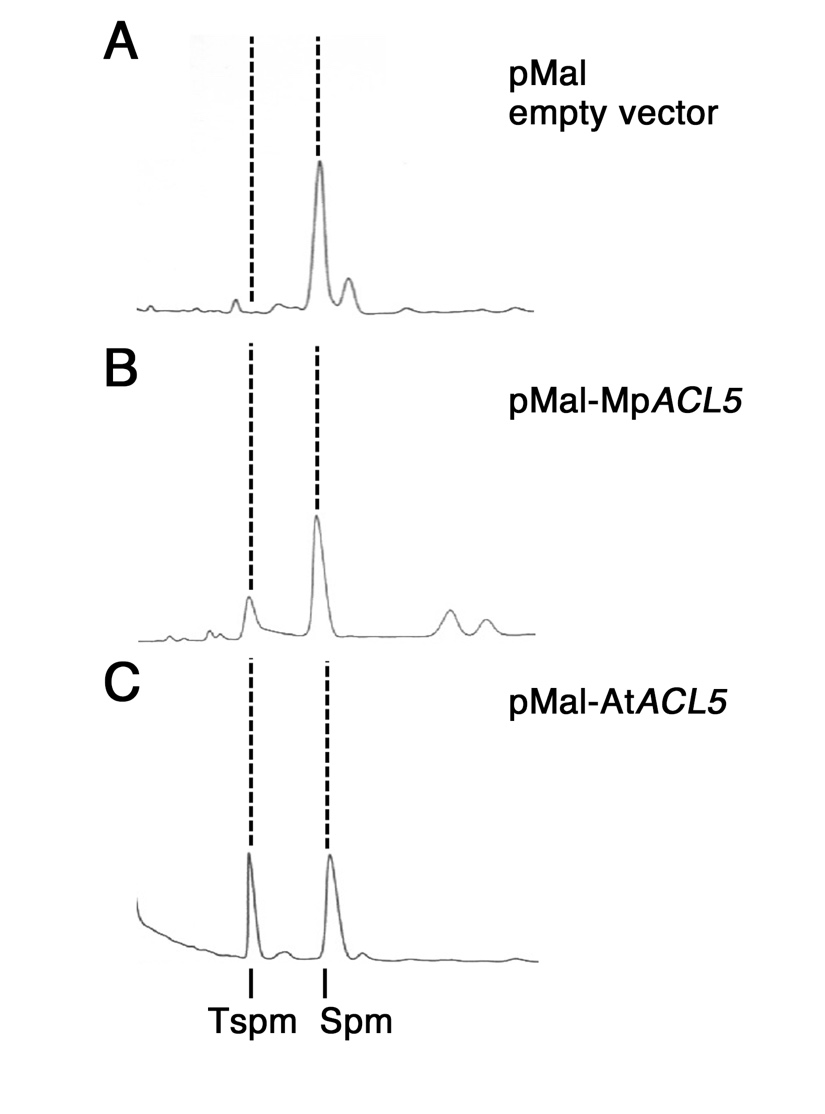

**Supplementary Fig. S2**	Bacterial expression of Mp*ACL5*. Polyamines were extracted from *E. coli* with the empty pMal vector, pMal-Mp*ACL5*, or pMal-At*ACL5* and subjected to the HPLC analysis after benzoylation to distinguish thermospermine and spermine. Spermine detected in all samples may be derived from culture media.

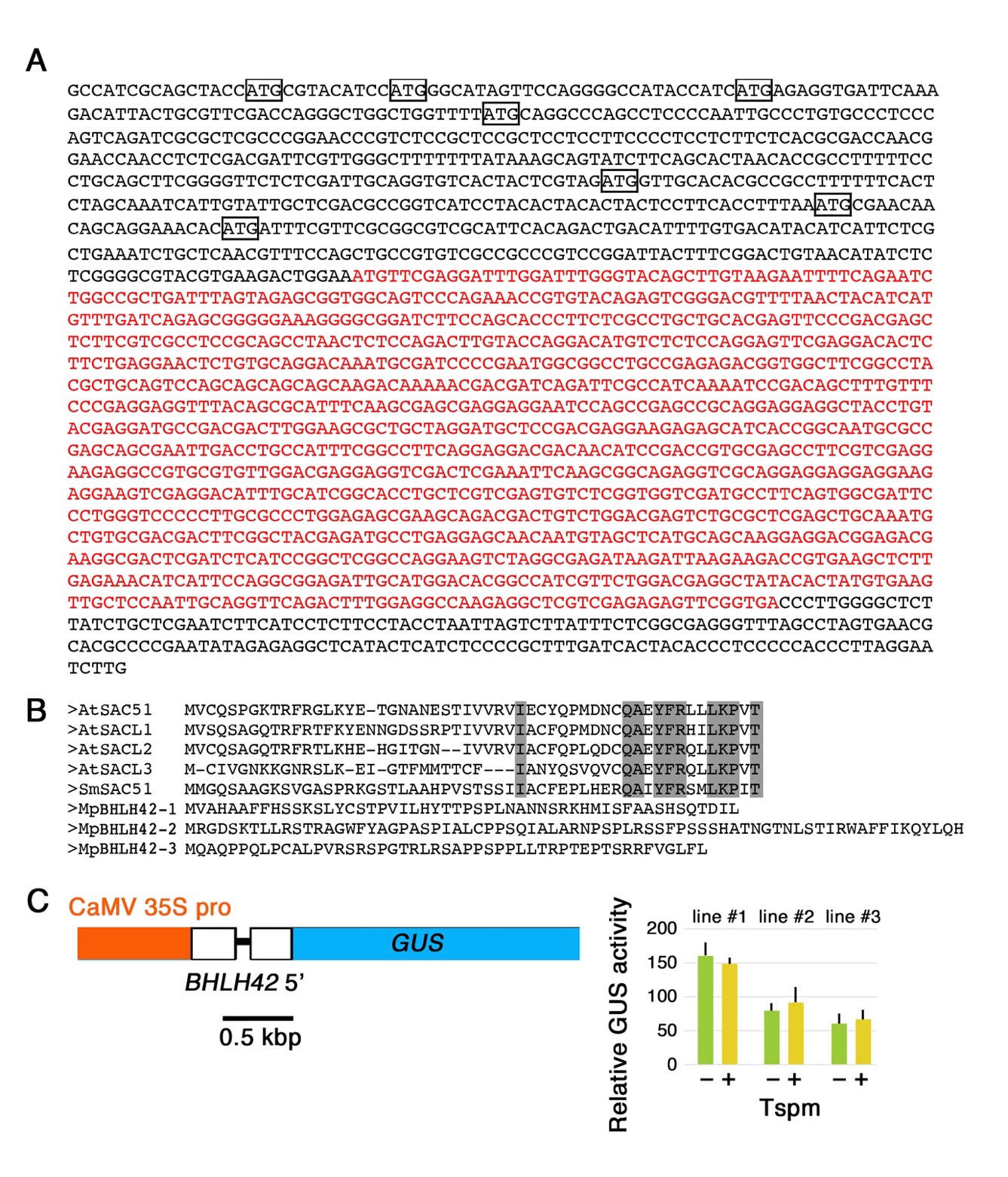

**Supplementary Fig. S3**	The closest homolog of *SAC51* in *M. polymorpha*, Mp*BHLH42***,** is not responsive to thermospermine. (A) The full-length cDNA sequence of Mp*BHLH42*. Start codons in the 5’ leader region are boxed. Main coding sequence is shown in red letters. (B) Alignment of peptide sequences encoded by uORFs conserved in the *SAC51* family in *Arabidopsis* and Selaginella moellendorffii (SmSAC51) with those present in Mp*BHLH42*. (C) The Mp*BHLH42* 5’ leader-GUS fusion construct and the GUS activity in transgenic *Arabidopsis* seedlings carrying the construct. The 5’ leader region was amplified from genomic DNA by PCR with primers, FXba, TCT AGAGCCATCGCAGCTACCATGC, and RBam, GGAT CCCAAATCCAAATCCTCGAAC, and inserted as a XbaI-BamHI fragment into pBI121. Activities in three independent transgenic lines are shown. Ten-day-old seedlings were incubated with 0.1 mM thermospermine for 24 h. Bars indicate SD (n=5).
